# Functional Diversity of Reef Fishes in the Veracruz Reef System National Park, SW Gulf of Mexico

**DOI:** 10.64898/2026.10.02.756167

**Authors:** Jesús Ernesto Arias-González, Aura Aletse Morales-Aranda, Diana Morales-de-Anda, Guillermo Horta-Puga

## Abstract

Reef fish assemblages in the Veracruz Reef System National Park (VRSNP) are strongly structured along a pronounced north–south gradient, where environmental conditions, urban proximity, and protection levels interact to shape functional diversity and ecosystem functioning. This gradient creates contrasting ecological conditions between northern and southern reefs, providing a natural framework for evaluating how spatial variation influences resilience. Using a stationary visual census across ten latitudinally distributed reefs, we quantified functional entities based on six ecological traits and calculated functional richness, redundancy, over-redundancy, vulnerability, specialization, and originality. Two-way permutational ANOVA and redundancy analysis were used to detect spatial and temporal patterns and to link functional indices to environmental features. The southern sector displayed the most pronounced functional shifts. Nearshore southern reefs, strongly exposed to sedimentation, nutrient loading, and physical disturbance, exhibited reduced functional richness, fewer functional entities, and high functional vulnerability, indicating strong environmental filtering. Over time, these reefs shifted toward generalist-dominated assemblages with rising redundancy, suggesting functional convergence. In contrast, southern offshore reefs supported higher taxonomic and functional richness, buffered by greater habitat complexity and greater distance from shore, although they showed declining redundancy, likely reflecting sensitivity to regional climatic disturbances. In the northern sector, the gradient was moderated by effective protection, which reshaped functional configurations. Nearshore protected reefs such as Isla Sacrificios and Isla de Enmedio maintained high redundancy, low vulnerability, and high biomass of large-bodied fishes, resembling offshore systems rather than disturbed southern nearshore reefs. The northern offshore reef (Isla Verde) shared some of these attributes but exhibited temporal declines in richness, signaling emerging vulnerabilities along the gradient. Functional originality and specialization aligned with benthic states across the gradient, particularly where macroalgal or calcareous substrates prevailed. Overall, the north–south gradient is the primary axis of functional organization in VRSNP, revealing that resilience emerges from the interplay among reef geomorphology, urban influence, and governance effectiveness.

## 1. Introduction

Understanding functional diversity, a subset of biodiversity, is essential for evaluating ecosystem health, resilience, and ecosystem services because it plays a critical role in ecosystem functioning [1–3]. Functional diversity refers to the range and variety of functional traits within a community assembly [4–7]. It emphasizes species’ roles in ecosystems rather than their abundance or richness [1,6,7]. Functional traits are specific attributes of a community assembly that influence ecological roles and interactions within that community [5,6]. Diverse functional traits can contribute to greater productivity, stability, and resilience in ecosystems, allowing them to withstand disturbances and recover more effectively [8,9]. They can enable species to exploit various resources, reducing competition and enhancing overall productivity [9]. They can also contribute to nutrient cycling in distinct ways, such as through variations in feeding habits or reproductive strategies, which, in turn, influence sediment and water quality and food webs [1,10].

Trait diversity encompasses several key components, including morphological, reproductive, and behavioral traits among species within a community [8,11,12]. For example, in a fish community, functional traits may include body size, feeding strategy, and habitat use. Species can be grouped by shared functional characteristics. For instance, herbivorous or piscivorous fish might be grouped because they perform similar roles, such as grazing on algae or feeding on fish. Multiple species may fulfill similar functional roles in the fish community, exhibiting functional redundancy [5]. High redundancy may enhance ecosystem resilience, as the loss of one species may not significantly affect ecosystem functions if others can fill that role. Conversely, the extent to which a single species represents specific functional roles indicates greater vulnerability and functional loss [5].

Analyzing spatiotemporal patterns of fish community functional traits provides critical insight into how ecological roles shift across reef locations and over time, revealing the dynamic interplay between environmental pressures and biological responses that shape ecosystem resilience [13,14]. These patterns capture how the functional characteristics of fish populations vary across spatial scales and over time [6]. Functional diversity and functional trait space offer powerful frameworks for understanding ecological patterns and detecting changes in fish community structure [15,16]. Temporal studies of fish functional traits can highlight patterns associated with ecological changes, species interactions, and human impacts [17]. This is crucial for understanding ecological dynamics, biodiversity, and the resilience of coral reef ecosystems [18]. Understanding the distribution of functional traits within ecological communities is key to elucidating the mechanisms that govern community assembly and persistence. However, despite growing recognition of its importance, spatiotemporal variation in the functional structure of marine fish communities (particularly within biodiverse and dynamic systems such as the Veracruz Reef System National Park) remains poorly studied [19].

The urgent need to predict the biological consequences of rapid ecosystem changes driven by human activities requires a clear understanding of ecological relationships [13]. The Veracruz Reef System (SAV), located in the southwestern Gulf of Mexico, has been exploited by humans since the arrival of the Spanish. Today, the Veracruz Reef System is part of the Veracruz Reef System National Park (VRSNP), which encompasses several reefs that face various environmental stressors. The VRSNP was established to conserve diverse aquatic ecosystems, including coral reefs, mangroves, and coastal lagoons. It plays a vital role in protecting marine biodiversity by providing habitat for fish species, invertebrates, and sea turtles. The VRSNP also supports local fisheries and tourism, encouraging the sustainable use of its resources. Conservation efforts in the VRSNP focus on monitoring reef health, managing human activities, and raising awareness of marine conservation.

The most significant stressors are the discharge from the Jamapa River and anthropogenic pressures associated with port activities [20,21]. Based on community structure and conservation status [21], three reef clusters were identified: nearshore or degraded, offshore or moderately conserved, and conserved. The entire Veracruz Reef System is regarded as moderately conserved. Previous studies on fish density indicated that it is highly correlated with the cover of hermatypic coral and crustose coralline algae [21]. However, whether these associations maintain consistent patterns remains unclear, as does whether specific reefs harbour fish with distinct functional traits. A study conducted between May 2006 and April 2021 examined the functional diversity of reef fish across seven different reefs of the VRSNP [22]. The research considered both southern and northern reefs, as well as inner- and outer-shelf reefs at both shallow and deep depths. It employed functional richness and divergence indices to show that defining geographical subsystems at specific depths where traits diverge would be more effective than defining them by north-south geographic location.

Our study aims to analyze the functional diversity of reef fishes in the VRSNP by evaluating their spatiotemporal functional structure. We seek to identify the factors that drive variability in this structure. Our goals are to: 1) Identify the functional structure of reef fishes in the VRSNP using a functional trait approach; 2) Determine the functional redundancy and vulnerability of reef fishes; 3) Explore differences in the assemblage composition of reef fishes across sites, seasons, and years, with a focus on large piscivorous, herbivorous, and planktivorous species that are key in the VRSNP; and 4) Determine whether coral or macroalgae cover and reef area play a role in shaping changes in functional diversity traits, vulnerability, and redundancy.

Our work hypothesized that fish functional traits contribute more to resilience and vulnerability than species richness alone. This effect is particularly evident on reefs with higher coral cover, larger island sizes, greater distance from continental landmasses, and formal protection status, where macroalgal dominance is reduced and anthropogenic pressures are minimized. This establishes reef size and distance from human stress as among the most consistent and essential drivers of functional diversity traits and resilience, with patterns emerging across spatiotemporal scales. Consequently, a more precise approach is to define sites, seasons, and years using specific variables that capture trait variation. Management and protection policies should focus on conserving areas with high trait diversity, redundancy, and low vulnerability, while also restoring biodiversity in sites with high vulnerability, low redundancy, and limited functional diversity.

## 2. Materials and Methods

### 2.1. Study Area

The study was conducted in the Veracruz Reef System National Park (VRSNP) in the southwestern Gulf of Mexico (Fig. 1). The VRSNP receives freshwater input from the Jamapa River, which reduces visibility due to suspended particles and elevates nutrient levels [21,23]. Climate, oceanography, and continental runoff shape the area’s seasonality, which comprises three climatic periods: the rainy season (June to October), the season of northern cold fronts (October to April), and the dry season (March to June) [20,24,25]. The VRSNP includes various reef formations that differ in habitat features, size, and levels of degradation [21]. The reefs selected for this study were classified by conservation status using the ratio of reef builders to macroalgae (RB/FM), as defined by Horta-Puga et al. [21].

1. Nearshore reefs dominated by non-reef-building communities (Hornos, Galleguilla, Isla Sacrificios, and Pajaros) were classified as degraded. Northern Reefs.
2. Offshore reefs (Anegada de Afuera, Blanca, Cabezo, and Isla Verde) were considered moderately conserved. Southern Reefs.
3. Reefs with benthic reef-building communities (Chopas and Isla de Enmedio) were classified as conserved. Southern Reefs.

**Fig. 1.**
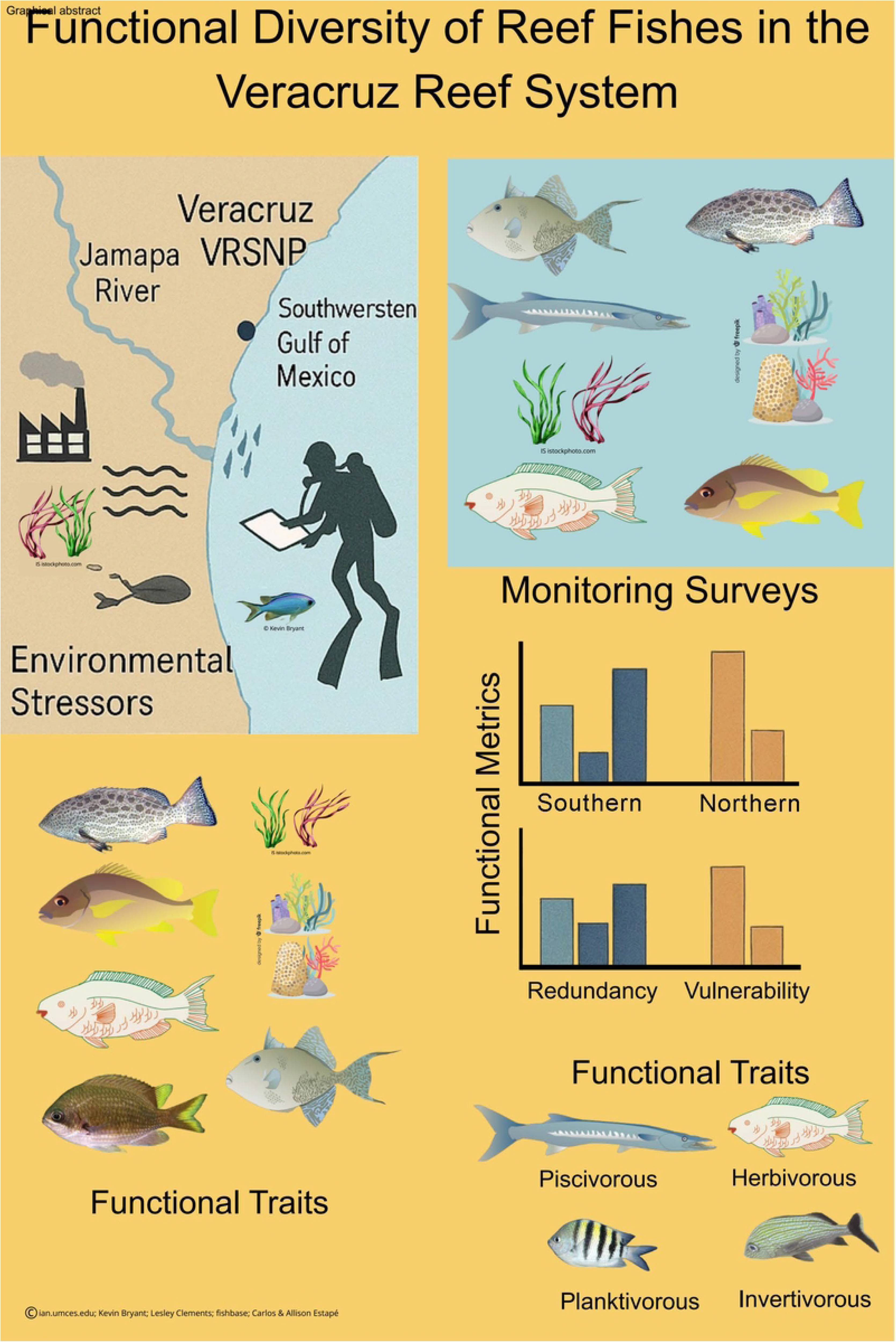
The Veracruz Reef System National Park (VRSNP) and the location of reef sampling sites.

### 2.2. Data Collection

Data were collected through five monitoring surveys carried out from 2012 to 2015 on ten reefs within the VRSNP (Fig. 1). To account for temporal variability, surveys took place during different seasons (dry: March/April; rainy: September/October): October 2012, March 2013, September 2013, March 2014, and March 2015. These surveys form part of a long-term census of the VRSNP [22].

A modified version of the stationary visual census method, as described by Bohnsack and Bannerot [26] and Horta-Puga et al. [23], was used to record fish species and abundance. This adaptation addressed the area’s low visibility, which was caused by freshwater discharge from the Jamapa and Papaloapan rivers [27]. Two divers performed six censuses per site (three per dive), recording conspicuous reef fish species within a 3 m radius and 6 m vertical column over 5 minutes, between 10:00 and 13:00 h.

#### 2.2.1. Benthic Macroalgae and Hermatypic Corals

Each sampling station was positioned on the leeward slope at 9–12 m depth to characterize the fish assemblage and benthic components. Macroalgae were categorized into three morpho-functional groups, frondose, turf, and crustose corallines, following Littler and Littler [28], and as implemented by Horta-Puga et al. [21]. An underwater camera mounted on a PVC frame with a 25 × 25 cm quadrat (625 cm² area) was used to photograph the substrate. Thirteen thousand six hundred eighty-eight photo-quadrats were analyzed, covering approximately 855 m² of reef. Calibrated observers on high-resolution displays estimated macroalgae cover. Coral cover (%) was assessed using live coral interception length along 10-m line transects until ≥100 coral colonies were sampled per site [25].

#### 2.2.2. Fish Traits

To assess the functional ecology of reef fish, we used six categorical functional traits widely applied in similar studies [5]: total length interval, trophic group, mobility, water column position, schooling behavior, and diel activity (Table S1). Ecological functional metrics and indices, along with functional trait data, were used to group species with the same trait values into Functional Entities (FEs). With this matrix, we calculated three ecological metrics [5]: functional redundancy (FRed), the number of species sharing the same trait combination; functional over-redundancy (FOver), the degree to which some FEs are overrepresented in terms of species richness; and functional vulnerability (FVul), the proportion of FEs represented by only one species. Additionally, we calculated functional diversity indices based on a functional space constructed using Gower distances and a Principal Coordinates Analysis (PCoA) of all fish functional entities (FEs). We assessed the number of axes required for the PCoA to provide a more accurate representation of the distances and trait differences between FEs, following the steps outlined in Maire et al. [29]. Based on this functional space, we calculated three additional and complementary multi-trait functional diversity indices [5]: 1) Functional richness

(FRic): the volume of multidimensional trait space occupied by the community. 2) Functional specialization (FSpe): the degree to which species have extreme trait values or are ecological specialists [30]. 3) Functional originality (FOri): the uniqueness of each species’ traits within the assemblage [31].

### 2.3. Statistical Analysis

We evaluated differences in metrics and indices across reefs and censuses using two-way permutational ANOVA (PERMANOVA) with 9,999 permutations and a Euclidean distance matrix [32]. To assess shifts in species composition by size and diet across sites and seasons, we used PERMANOVA with a Bray–Curtis similarity matrix and a one-factor model. Similarity Percentage Analysis (SIMPER) identified the functional entities that contributed most to the observed differences. To visualize shifts in fish community composition, we applied non-metric multidimensional scaling (NMDS) and highlighted changes in biomass among large-bodied and piscivorous species. All analyses were performed using PRIMER v6 and PERMANOVA+ [33,34].

## 3. Results

### 3.1 North–South Patterns in Functional Structure and Diversity

Clear latitudinal differences (south vs. north reefs, Fig. 1) were evident in the functional structure, diversity, and trophic organization of reef fish assemblages in the Veracruz Reef System National Park (VRSNP). These differences were modulated by reef position relative to the coast, the degree of urban influence, and protection status.

Across the park, 116 reef fish species were grouped into 89 Functional Entities (FEs), with most FEs represented by a single species, indicating high functional vulnerability (Fig. 2a; Table S1). Invertivorous fishes dominated the functional space in both regions, followed by piscivorous, herbivorous, and planktivorous groups. Functionally vulnerable FEs were consistently located at the margins of the functional trait space (Fig. 2b), especially on reefs exposed to anthropogenic disturbance (Fig. 1).

**Fig. 2.**
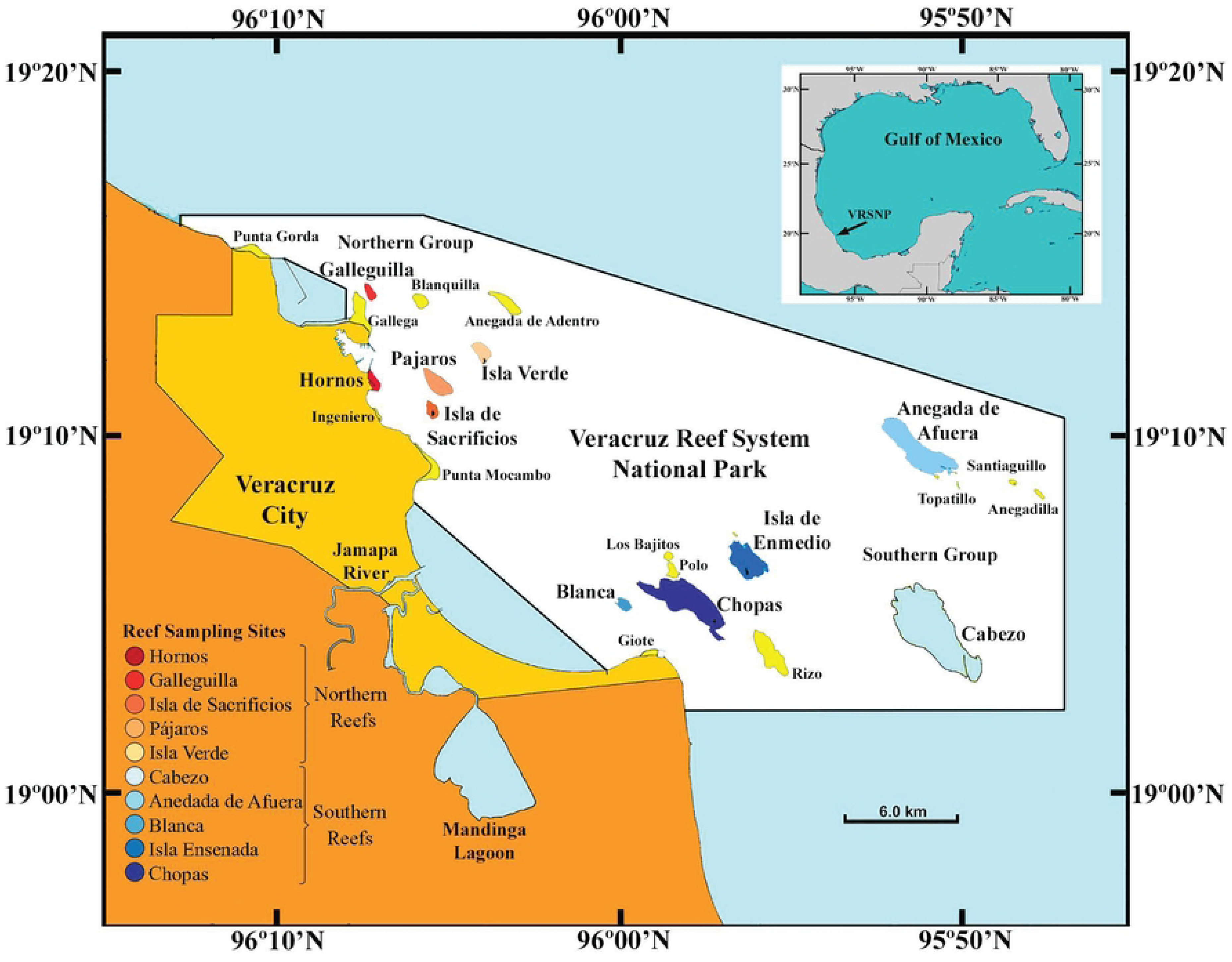
General representation of fish species and traits (six traits) found across the 10 reefs from the VRSNP. (a) Bar plot displaying the number of species per Functional Entity (FE) categorized by diet. (b) Principal Coordinates Analysis (PCoA) of FEs using Gowers distance based on functional traits; the circle denotes the FE in functional space, the number within the circles highlights those FEs at the edge of the functional space, and this number corresponds to the specific FE number matching those in the (a) bar plot. The colors in the bars (a) and points (b) indicate the diet trait, one of the six traits used to describe reef fish.

### 3.2. FEs, Fred, FOver and FVul across Reefs and Censuses

We have observed significant variability across space (Reefs), time (Census), and their interaction (Reefs × Census). Considerable variability in FEs, Fred, FOver, and Fvul values was evident in the highly significant differences observed in the two-way crossed (Census × Reefs) permutational ANOVA results (Table 1).

**Table 1.**
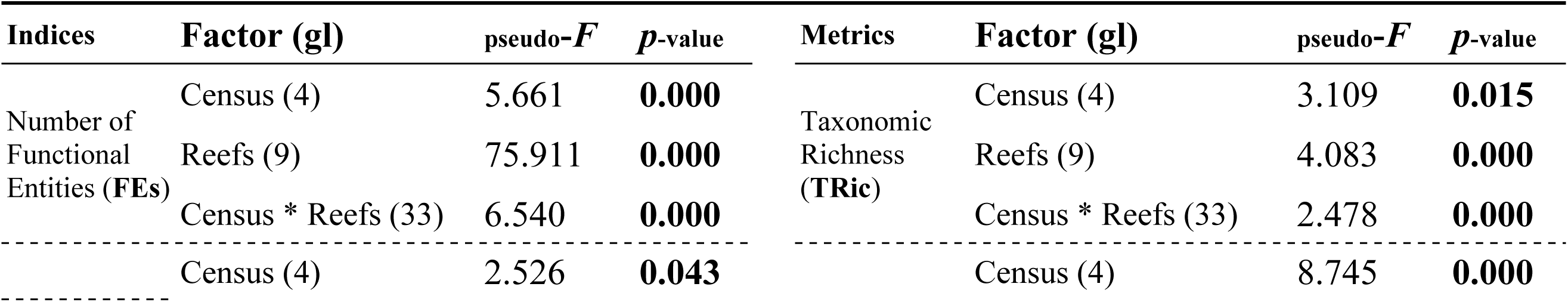
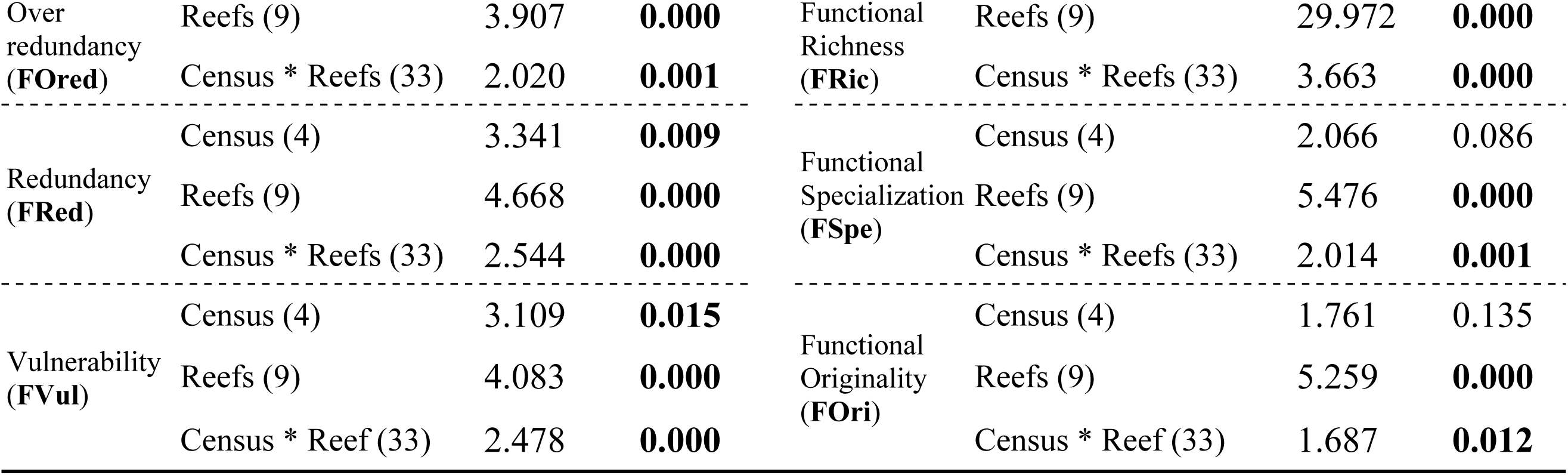
Two-way crossed (Census x Reefs) permutational ANOVA results (9999 permutations) for functional metrics: FEs, FOred, FRed, FVul, and indices: TRic, FRic, FSpe, and FOri for the Veracruz Reef System National Park (VRSNP)

### 3.2. Southern Reefs

Southern reefs exhibited the strongest spatial contrasts in functional organization. Offshore southern reefs, notably Cabezo and Anegada de Afuera, supported higher numbers of FEs (Fig. 3a) and lower functional vulnerability (Fvul; Fig. 3d), greater taxonomic richness (Tric, Fig. 4a), and moderate-to-high functional richness (Fric, Fig. 4b), These reefs are large, structurally complex, and distant from urban development, providing favorable conditions for functionally diverse assemblages (Fig 3a-d; Fig. 4a-d).

**Fig. 3.**
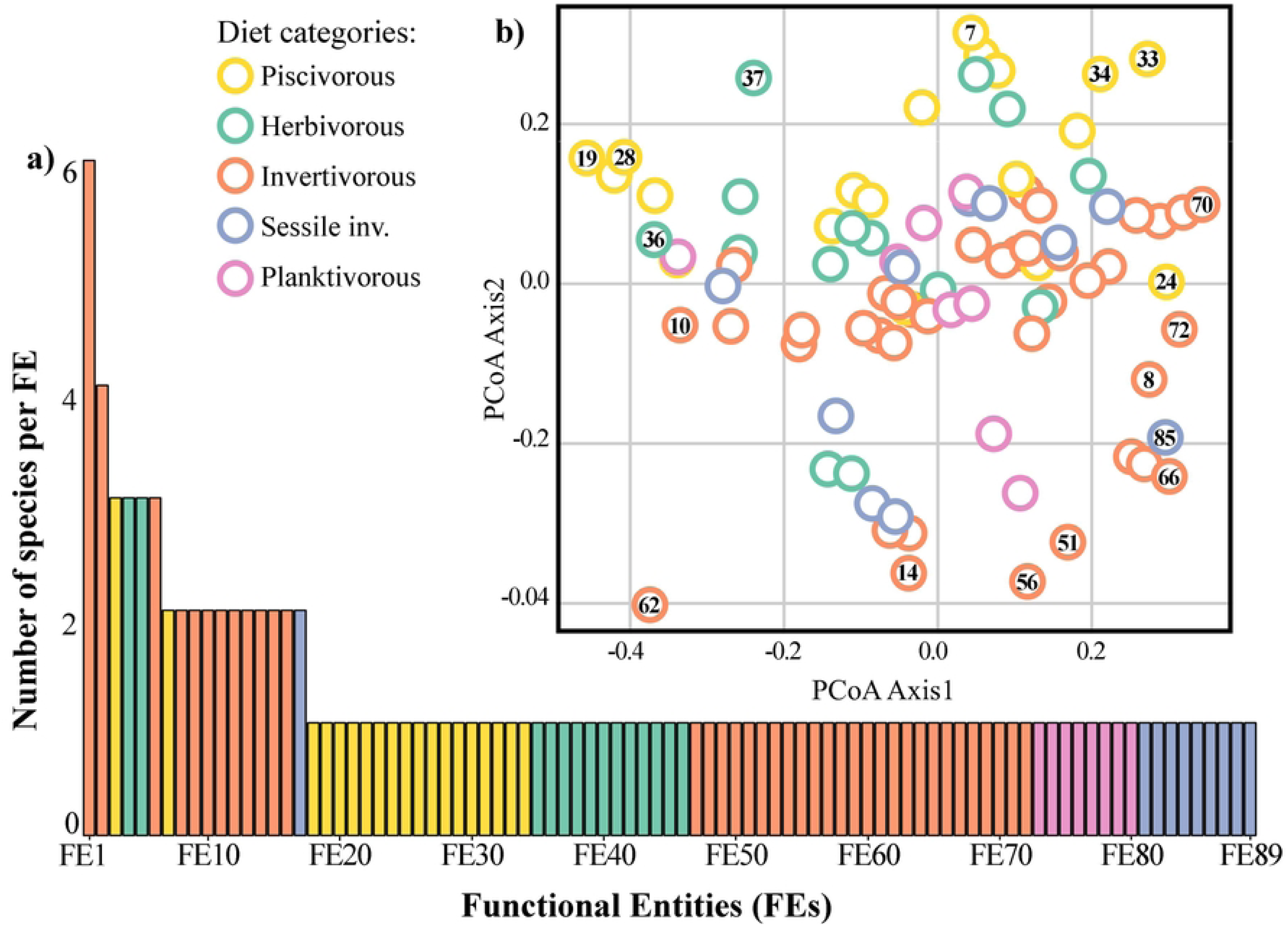
Functional metrics across different Reefs in VRSNP. Average values with standard deviation (±SD) are shown for temporal variation across censuses: (a) Number of Functional Entities (FEs), (b) Functional Redundancy, (c) Functional Over-redundancy, and (d) Functional Vulnerability. Each point in all graphs represents a reef site, with colors consistently representing the same reef across all graphs.

**Fig. 4.**
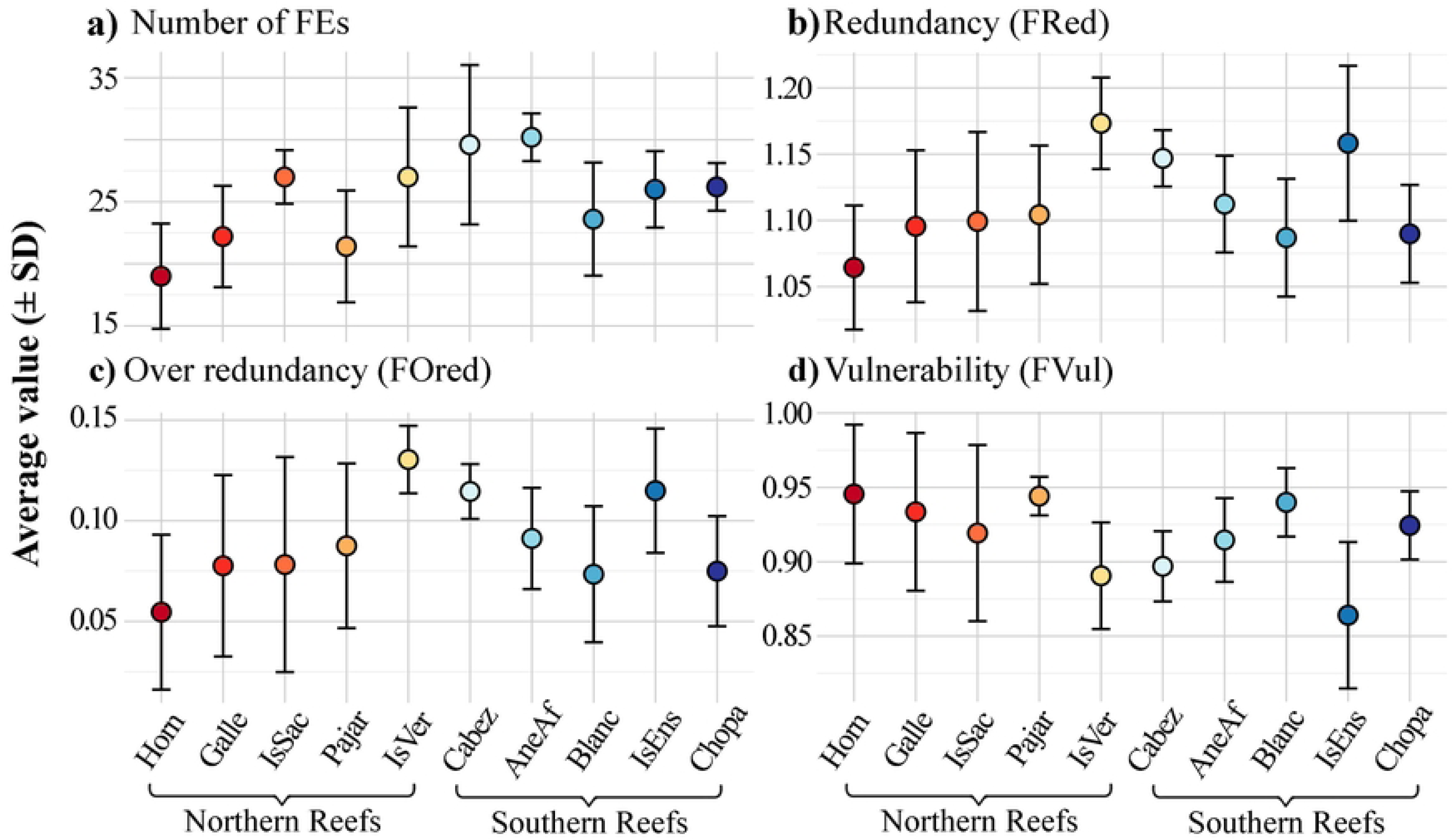
Diversity indices across different reefs in VRSNP. Average values with standard deviation (±SD) are shown for temporal variation across censuses: (a) Taxonomic richness (TRic), (b) Functional richness (FRic), (c) Functional specialization (FSpe), and (d) Functional originality (FOri). Each point in all graphs represents a reef site, with colors consistently representing the same reef across all graphs.

In contrast, nearshore southern reefs (Hornos, Galleguilla, and Pájaros) showed fewer FEs, lower redundancy (FRed and FOred), and higher vulnerability (Fig. 3a-d). These reefs also displayed consistently high functional originality (FOri), indicating assemblages dominated by a limited number of functionally unique species (Fig. 4d).

Temporal trends showed declining FEs in Hornos, Galleguilla, and Pájaros, whereas offshore southern reefs gained FEs over time but simultaneously lost redundancy (Fig. S2). Functional vulnerability decreased over time in the most urban-influenced southern reefs, suggesting a shift toward dominance by generalist species with overlapping functions (Fig. S2).

### 3.3. Northern Reefs

Northern reefs showed less pronounced latitudinal gradients, yet distinct site-level patterns emerged. Isla Sacrificios, despite being nearshore, exhibited high FRed and FOred and low functional vulnerability (Fig 3a-d). This reef also supported high biomass across trophic groups and size classes, likely reflecting its protected status and restricted public access. Isla de Enmedio showed similar patterns, underscoring the importance of protection in mitigating nearshore impacts. In contrast, Isla Verde, an offshore northern reef, displayed high TRic and moderate FRic, resembling offshore southern reefs in functional structure (Fig. 4 a-d).

Temporal variability in the north included declines in TRic and FRic at Isla Verde and Isla Sacrificios toward the end of the study period, accompanied by reef-specific shifts in functional specialization (FSpe) and originality (Fig. 2S).

### 3.4. Functional Diversity and Environmental Drivers

Functional richness peaked at reefs at intermediate distances from urban development in both sectors (e.g., Chopas in the south and Isla Sacrificios in the north) (Fig. 4b). Functional specialization showed limited spatial structure (Fig. 4c), except for high temporal variability at Galleguilla (Fig. 2S). Functional originality was elevated in both disturbed nearshore reefs and select offshore reefs, suggesting alternative pathways to functional uniqueness (Fig. 4d).

Redundancy analysis revealed a consistent north–south structure associated with habitat characteristics. TRic and FRic were positively associated with offshore reefs characterized by high coral cover and larger reef area, whereas FSpe and FOri were linked to reefs dominated by macroalgae and calcareous algae, respectively (Fig. 5). These relationships were more tightly expressed in southern reefs.

**Fig. 5.**
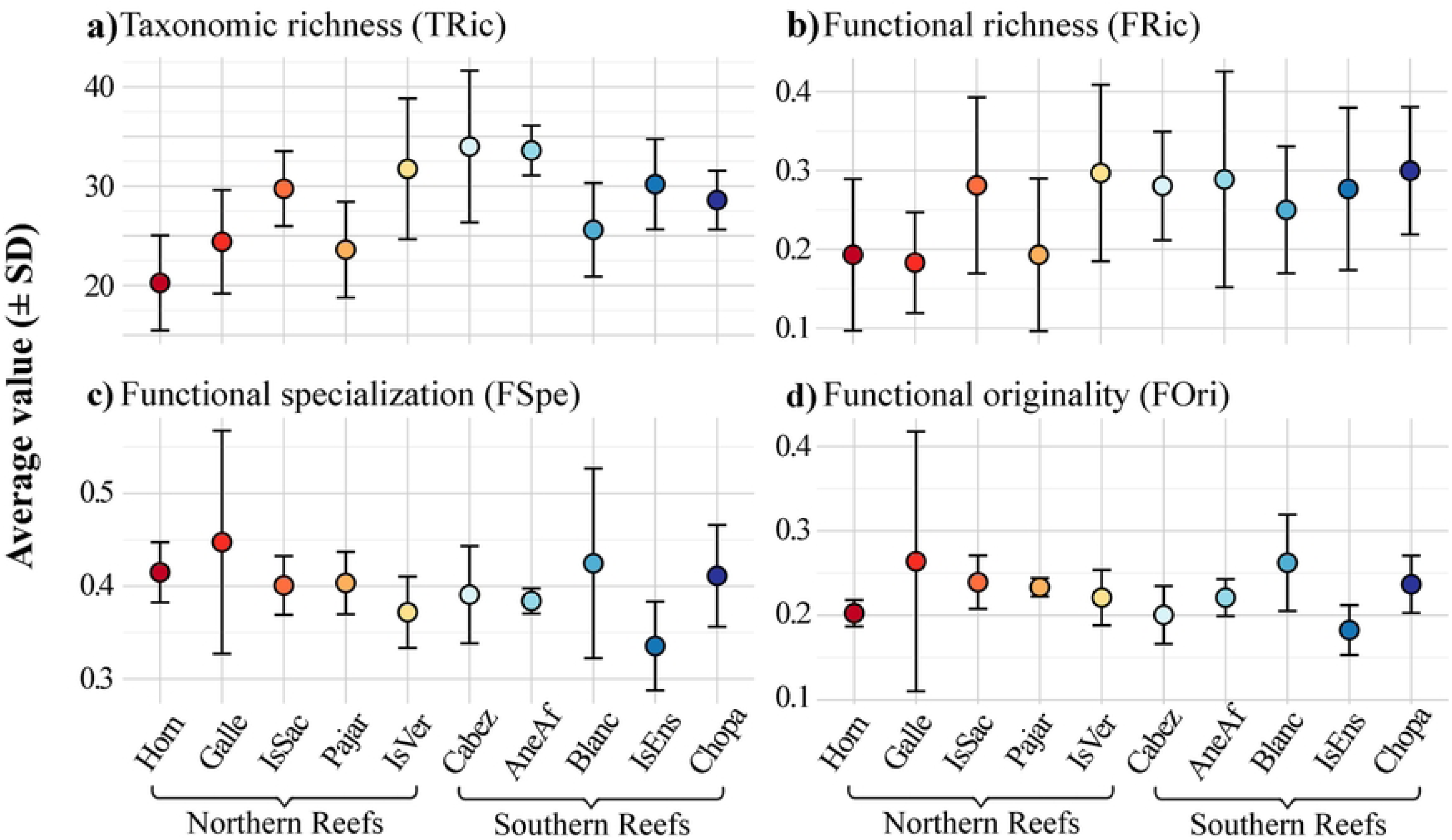
Redundancy Analysis (RDA) plot displaying the relationship between environmental variables and diversity indices. Points represent reef sites, with numbers indicating different censuses. Arrows denote the direction and strength of ecological gradients, and species are labelled in bold. The size and colour of the points correspond to the specific reef sites, with consistent colours used across censuses.

### 3.5. Trophic structure

Invertivorous fishes dominated biomass across both regions, followed by planktivores. Southern offshore reefs supported high biomass of herbivores and medium-sized fishes, while northern protected reefs (Isla Sacrificios and Isla de Enmedio) supported disproportionately high biomass of large-bodied fishes (Fig. 6). These latitudinal patterns highlight the combined influence of urban proximity, reef morphology, and protection status on functional diversity and ecosystem functioning in the VRSNP.

**Fig. 6.**
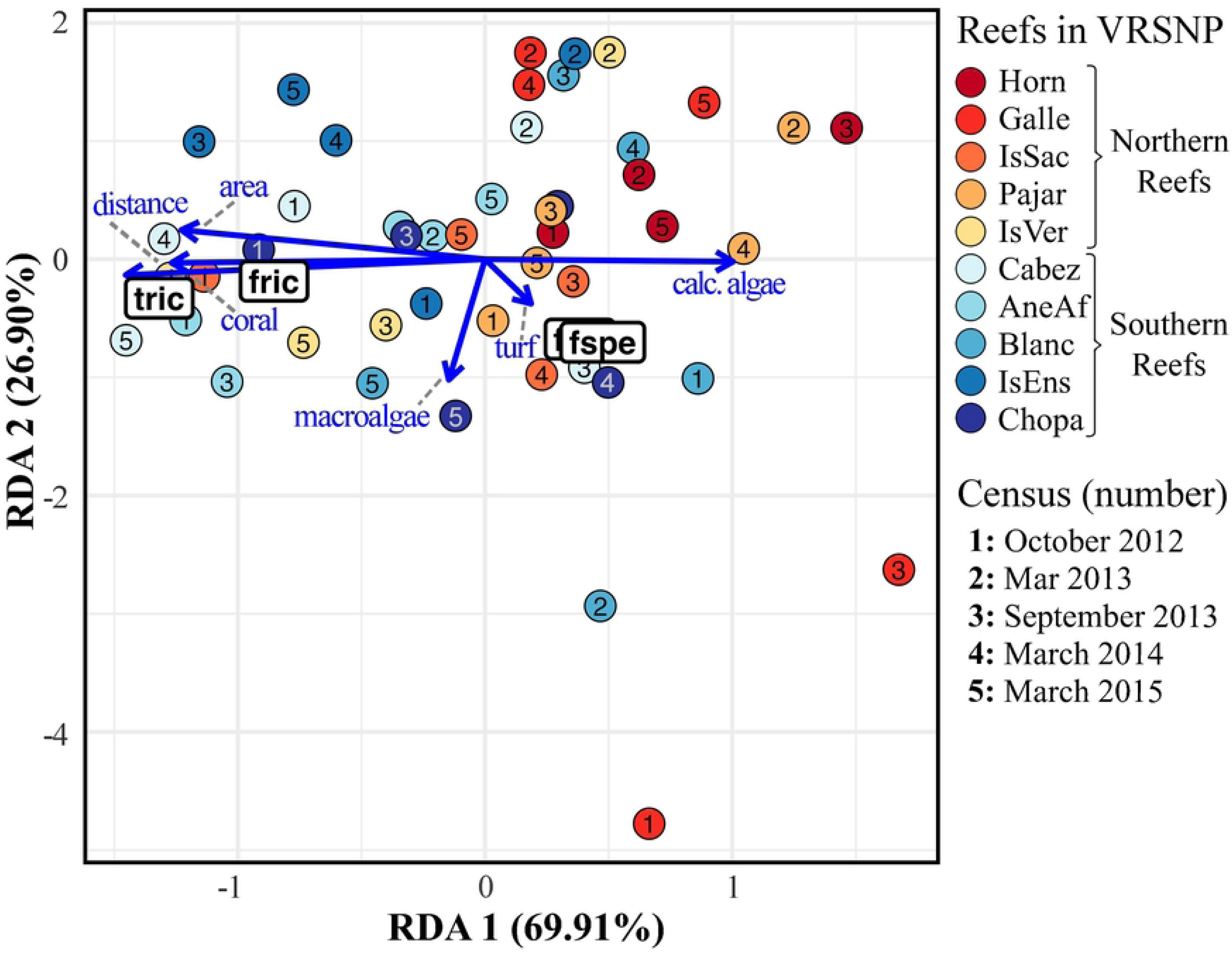
NMDS plot representing the reef fish community composition at VRSNP, the differences in composition of reef fish are represented by the bubbles, and the size of the bubbles indicates the average biomass per census per corresponding trait representation. Two traits categorize the plots: Left-side plots represent the diet trait, with (a) invertivorous (b) herbivorous (c) piscivorous fish and (d) planktivorous; the right-side plots represent the size trait, with (e) small (f) medium and (g) large fish.

**Fig. 7.**
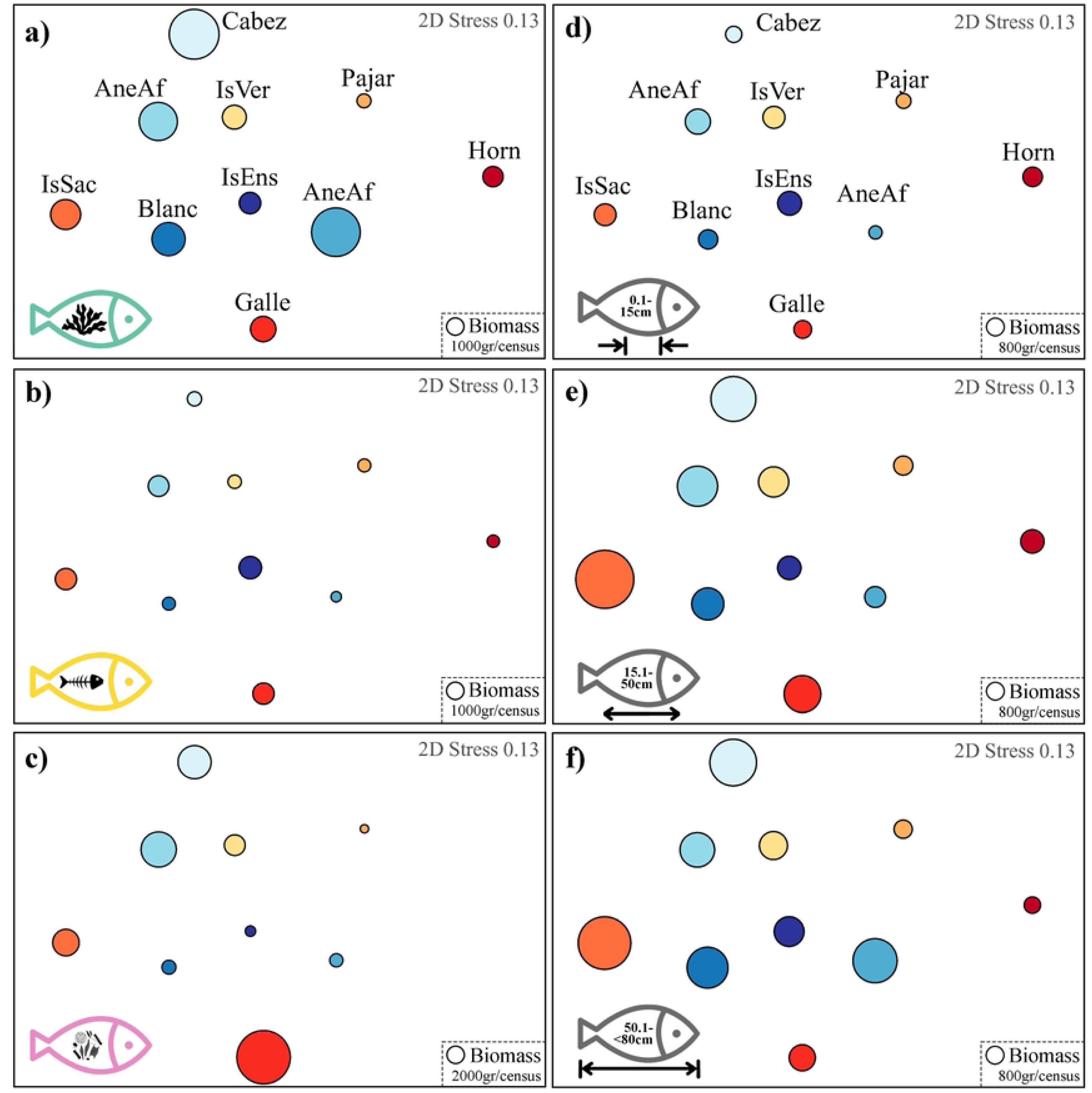

## 4. Discussion

The functional diversity and trophic structure of reef fish assemblages in the Veracruz Reef System National Park (VRSNP) exhibit a clear north–south structuring, reflecting the combined influence of urban proximity, reef geomorphology, habitat composition, and protection status [4,5]. While both sectors share a dominance of invertivorous fishes and a high proportion of functionally vulnerable entities, the mechanisms shaping redundancy, vulnerability, and trophic structure differ markedly between southern and northern reefs. Southern reefs showed the strongest functional gradients, particularly between nearshore, urban-proximal reefs and offshore, distal reefs. Nearshore southern reefs were characterized by lower functional richness and higher functional vulnerability, consistent with environmental filtering [35,36] from chronic exposure to anthropogenic stressors, including sedimentation, eutrophication, and physical disturbance [20,24,25,37,38]. These pressures disproportionately affect specialists and species with limited functional overlap, resulting in assemblages dominated by a narrower range of viable trait combinations [4,6,39,40].

Over time, functional vulnerability declined in these nearshore southern reefs, driven by increasing redundancy and over-redundancy [5]. This pattern suggests a replacement mechanism, whereby sensitive or functionally unique species are progressively replaced by generalist species with overlapping traits, producing a biotic homogenization [41,42]. While this may stabilize certain ecosystem processes in the short term, it is likely to reduce functional versatility and limit ecosystem resilience under future disturbances [43]. In contrast, offshore southern reefs maintained higher taxonomic and functional richness, reflecting the buffering role of reef size, structural complexity, and reduced land-based influence.

Northern reefs exhibited weaker latitudinal gradients, largely due to the presence of effectively protected nearshore reefs. Isla Sacrificios and Isla de Enmedio, despite their proximity to urban areas, displayed high functional redundancy, low vulnerability, and elevated biomass across trophic guilds and size classes [5]. These patterns highlight the critical role of access restriction and enforcement in mitigating fishing pressure and preserving functionally overlapping species that underpin ecosystem resilience [1,8,44,45]. The similarity in functional structure between protected northern nearshore reefs and southern offshore reefs underscores that management status can override spatial position. In contrast, unprotected northern reefs showed signs of declining taxonomic and functional richness toward the end of the study period, suggesting that regional stressors, potentially including thermal stress or episodic disturbances, may be increasingly influential even where local conditions are comparatively favorable [1,7].

Functional originality emerged as a key axis differentiating reefs between the two sectors. High originality was observed in heavily disturbed nearshore reefs and in select offshore reefs, indicating that functionally unique species persist under contrasting mechanisms [46]. In disturbed systems, high originality likely reflects assemblages of a few species occupying marginal or remnant functional niches, often with limited redundancy, thereby increasing vulnerability to further perturbation [31,46]. In offshore or protected reefs, high originality may instead reflect the maintenance of rare trait combinations supported by greater habitat heterogeneity and lower exploitation.

Functional specialization showed limited spatial structuring but strong temporal variability at certain reefs, particularly Galleguilla. Such variability suggests ongoing trait reorganization, potentially driven by fluctuating benthic conditions, shifts in resource availability, or transient disturbances [4,47,48]. Together, these indices indicate that changes in functional diversity are not solely directional but involve complex reconfiguration of trait space.

Redundancy analysis revealed consistent associations between functional indices and benthic habitat characteristics across both sectors. Reefs with higher coral cover and larger area supported greater taxonomic and functional richness, reinforcing the role of structural complexity in expanding habitat availability [1,8,44,45]. In contrast, macroalgae-dominated reefs were associated with higher functional specialization, whereas calcareous algae were associated with greater functional originality. These relationships suggest that benthic regime shifts directly translate into altered functional configurations of fish assemblages, with implications for ecosystem processes such as herbivory, bioerosion, and energy transfer [49–52].

Across the VRSNP, invertivorous fishes dominated biomass, underscoring the central role of benthic invertebrate pathways in reef trophodynamics [5,53]. However, trophic guild and body-size distributions varied among sectors. Southern offshore reefs supported high herbivorous biomass and medium-sized fish biomass, consistent with lower exploitation pressure and extensive grazing areas. Northern protected reefs supported disproportionately high biomass of large-bodied fishes, indicative of reduced size-selective fishing mortality and the persistence of upper trophic levels [22].

Maintaining large-bodied fishes at protected nearshore reefs likely enhances top-down control and stabilizes food-web interactions [54,55], thereby promoting functional redundancy and reducing vulnerability. These findings reinforce the importance of protection in sustaining not only species richness but also functional and trophic completeness [56,57].

## Conclusion

Together, the north–south patterns observed in the VRSNP show that functional resilience arises from the interplay of environmental gradients and governance. Offshore reefs naturally support higher functional diversity due to their size and isolation, whereas nearshore reefs can achieve comparable functional integrity when effective protection limits anthropogenic pressure. Conversely, urban-proximal reefs without adequate management converge toward functionally redundant, simplified assemblages, potentially masking underlying vulnerability.

From a management perspective, maintaining functional diversity in the VRSNP will require integrated strategies that combine land-based impact mitigation, targeted protection of nearshore reefs, and monitoring of functional redundancy trajectories. Such approaches are essential for preserving ecosystem functioning amid increasing regional and global stressors and for ensuring that functional diversity continues to buffer reef ecosystems against future change.

## CRediT authorship contribution statement

JEAG-DMDA-Conceptualization; GHP-AAMA-Data curation; DMDA-JEAG-Formal analysis; GHP-Funding adquisition; JEAG-DMDA-GHP-AAMA-Investigation; GHP-AAMA-Methodology; JEAG-DMDA-Writing-Original Draft; JEAG-DMDA-GHP-AAMA-Writing-review & editing. Supervision.

## Acknowledgements

The authorities of the Veracruz Reef System National Park (CONANP) granted the permits for this study. Manuel Victoria Muguira (Dorado Buceo) and Joseph Loustalot Laclette (Scubaver), helped us with the scuba dive gear. We appreciated the collaboration of Jose Luis Tello Musi, Alejandro Córdova, and Jhoan Gutiérrez Martinez (FES Iztacala, UNAM), during the field work.

## Data availability

Data will be made available on request.

